# DeFrND: detergent-free reconstitution into native nanodiscs with designer membrane scaffold peptides

**DOI:** 10.1101/2024.11.07.622281

**Authors:** Qian Ren, Jing Wang, Vinay Idikuda, Shanwen Zhang, Jeehae Shin, W. Grant Ludlam, Luis M Real Hernandez, Ilya Levental, Kirill Martemyanov, Baron Chanda, Huan Bao

## Abstract

Membrane scaffold proteins-based nanodiscs (NDs) have facilitated unprecedented structural and biophysical analysis of membrane proteins in a near-native lipid environment. However, successful reconstitution of membrane proteins in NDs requires prior solubilization and purification in detergents, which may impact their physiological structure and function. Furthermore, the detergent-mediated reconstitution of NDs is unlikely to recapitulate the precise composition or asymmetry of native membranes. To circumvent this fundamental limitation of traditional ND technology, we herein describe the development of membrane-solubilizing peptides to directly extract membrane proteins from native cell membranes into nanoscale discoids. By systematically protein engineering and screening, we created a new class of chemically modified Apolipoprotein-A1 mimetic peptides to enable the formation of detergent-free NDs (DeFrNDs) with high efficiency. NDs generated with these engineered membrane scaffold peptides are suitable for obtaining high-resolution structures using single-particle cryo-EM with native lipids. To further highlight the versatility of DeFrNDs, we directly extract a sampling of membrane signaling proteins with their surrounding native membranes for biochemical and biophysical interrogations.

## Introduction

The invention of nanodiscs (NDs) by the Sligar group twenty years ago revolutionized all aspects of membrane biology^1, 2^. By engineering amphipathic membrane scaffold proteins (MSPs) from Apolipoprotein-A1 (AopA-I), membrane protein complexes can be embedded in a lipidic environment for in-depth biophysical characterizations that are usually challenging in traditional membrane mimetic systems. Although the ND technology has emerged as a powerful tool for various applications, its assembly principle is rather straightforward^3^. The hydrophobic face of MSPs encircles a nanoscale patch of lipid bilayers inside NDs for the reconstitution of membrane proteins, whereas the outside hydrophilic face ensures water solubility of the assembled discoidal particles. Since the first generation of MSPs reported by Bayburt et al., the simple yet elegant framework of NDs inspired extensive efforts to further expand the structure and function of this highly useful platform using distinct designs of protein and polymer scaffolds^4–12^. These studies have culminated in an increasingly versatile ND toolkit for reconstituting diverse lipids and membrane proteins of interest. In addition, the rigid structure of NDs with diameters ranging from 5-50 nanometers has enormous therapeutic potential as a delivery vehicle for small molecules, vaccines, and immunotherapeutic drugs^13–20^.

Despite the tremendous progress in the development and application of ND technology over the past decade, many membrane protein complexes remain recalcitrant to this approach. The main difficulty is that current ND reconstitutions rely on the self-assembly of lipids, MSPs, and membrane proteins of interest upon the removal of detergents^21^. A prerequisite of this assembly process is to obtain stable membrane proteins in detergent micelles. Unfortunately, this prerequisite is a notoriously unmet challenge for membrane protein complexes that often unfold or fall apart once extracted from lipid bilayers using detergents. To address this problem, chemical crosslinkers are employed to increase the stability and homogeneity of membrane protein complexes for structural studies^22, 23^. However, these crosslinked protein complexes are no longer suitable for other biochemical and functional interrogations because their dynamics are markedly distorted. Therefore, a pressing need exists to bypass the limitation of detergent-mediated reconstitution of NDs.

On this front, amphipathic polymers have been developed for detergent-free reconstitution of membrane proteins into native NDs^7^. In recent years, these NDs have been widely utilized to elucidate the function of biological membranes in regulating the conformational state and stoichiometry of membrane proteins in diverse transmembrane signaling pathways^24–26^. However, the sensitivity of polymer NDs for bivalent cations and the altered packing of extracted native lipids makes it challenging for functional characterizations of many membrane proteins^12, 27–29^. Previous studies have also shown that screening a large library of different polymers for ND reconstitution is often required to overcome some unforeseeable difficulties. In addition, lipids undergo rapid exchange between these NDs^30^, indicating that the polymer-enclosed membranes will need further optimization to faithfully resemble the typical characteristics of lipid bilayers.

Herein, we propose to develop a different and complementary approach for detergent-free ND (DeFrND) reconstitution through the engineering of membrane scaffold peptides. The design of this approach is based on previous studies of membrane disruption by antimicrobial and ApoA-I mimetic peptides^13, 14, 31–36^. Through extensive rational design and screening of these amphipathic peptides, we generate a library of membrane scaffold peptides capable of directly extracting membrane protein complexes from native cell membranes into NDs without the need for detergents. The resulting NDs can maintain integral membrane proteins in functional states and provide an appropriate lipidic environment for the characterization of peripheral membrane proteins. Moreover, our approach allows for the preservation of membrane protein complexes that are usually disrupted by detergents, thereby enabling biophysical interrogations of detergent-sensitive membrane proteins unattainable in previous studies. Finally, we demonstrate that our detergent-free native ND technology using re-engineered membrane scaffold peptides is compatible with a wide range of transporters, receptors, and ion channels in bacterial and eukaryotic cells.

## Results

### Screening membrane scaffold peptides for detergent-free ND reconstitution

It is well established that amphipathic peptides can drastically remodel membranes and transform small unilamellar vesicles into discoidal ND structures without detergents^37^. By virtue of this advantage, these peptides are invaluable reagents for therapeutic development against bacterial resistance, cancer vaccines, and viral infections^14, 38, 39^. Inspired by these findings, we set out to characterize the ability of these peptides to extract a bacterial prototype ATP-binding cassette transporter MalFGK_2_ into NDs from proteoliposomes (Fig. 1a). MalFGK_2_ is composed of two transmembrane proteins (MalF and MalG) and two copies of the ATPase component (MalK) that powers the uptake of maltose into bacteria in the presence of the maltose binding protein (MBP)^40^. Previous studies have shown that the basal ATPase activity of MalFGK_2_ in proteoliposomes is low and tightly coupled to the transport of maltose^41, 42^. In contrast, the detergent-solubilized MalFGK_2_ complex is over 100-fold more active and is uncoupled from maltose and MBP. Thus, the ATPase activity of MalFGK_2_ is highly sensitive to the lipid environment and is a useful feature in evaluating the performance of membrane mimetic reconstitution systems.

**Figure 1.**
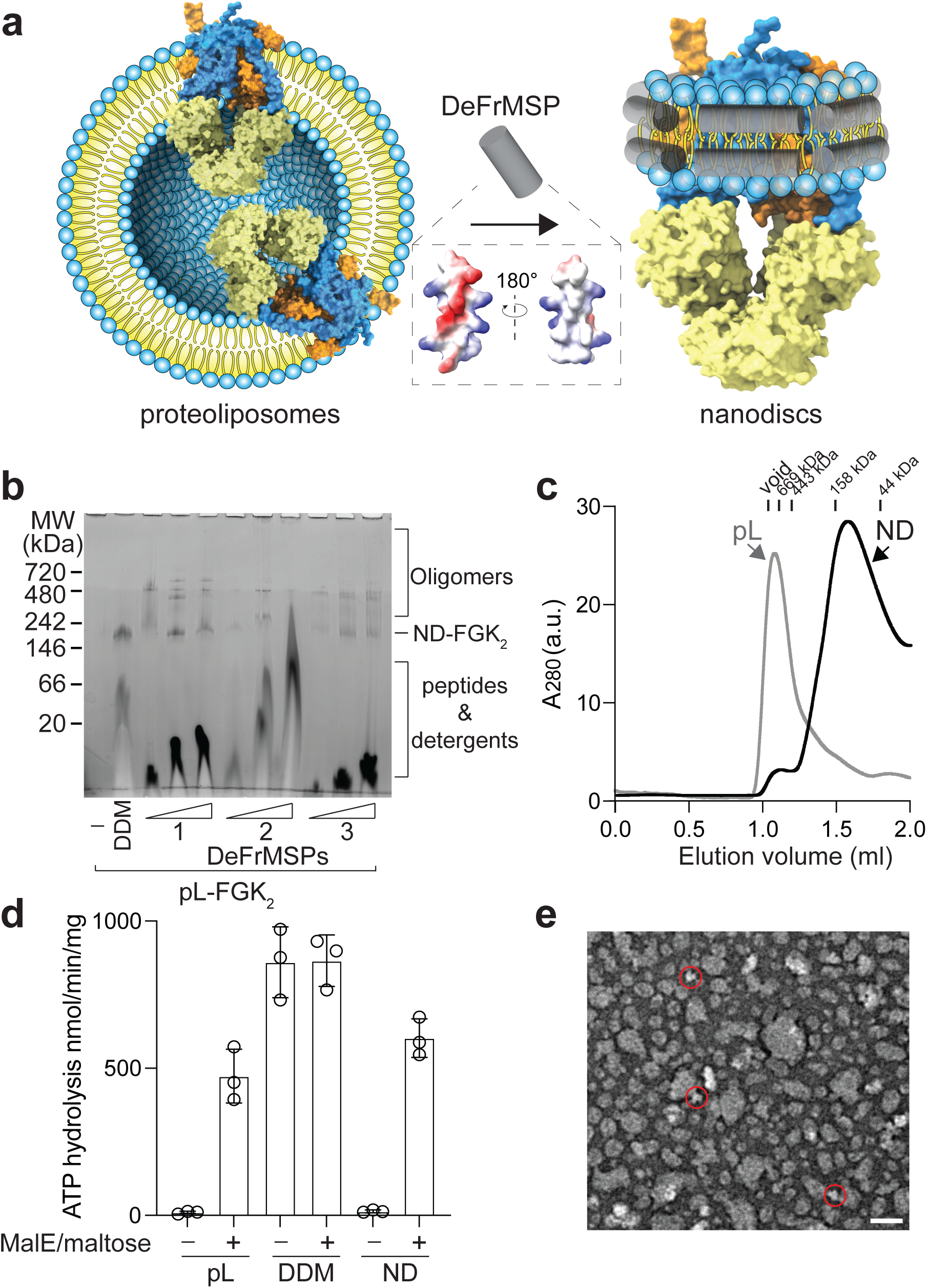
Screening membrane scaffold peptides for detergent-free reconstitution of NDs. (**a**) Illustration of the peptide screening strategy. Potent scaffold peptides will insert themselves into lipid bilayers of proteoliposomes and extract the maltose transporter MalFGK_2_ into NDs. Inset, surface charges of DeFr_1 as an exemplary amphipathic scaffold peptide. The structural model of of DeFr_1 is generated by Alphafold2. MalF, blue; MalG, orange; MalK, yellow. (**b**) Blue native gel analysis of detergent-free ND reconstitution. MalFGK_2_ proteoliposomes prepared using PC lipids were incubated with the indicated peptide scaffolds (33, 100 and 300 µM) in the absence of detergents and then analyzed by native electrophoresis. In control experiments, the same proteoliposomes were analyzed alone (lane 1) or treated with DDM (lane 2). (**c**) SEC profiles of detergent-free MalFGK_2_ NDs formed from proteoliposomes (pL) using of DeFr_1, in comparison with control experiments of pL alone. Samples were fractionated on a Superdex S200 3.2/300 column. (**d**) ATPase activities of MalFGK_2_ in detergent micelles, pL, and of DeFr_1-enclosed NDs. Data are shown as mean ± s.d., n = 3 independent experiments. (**e**) Negative stain EM micrograph of MalFGK_2_ NDs encased by of DeFr_1. Red circles highlight the monomer NDs. Scale bar, 30 nm.

We began our study with a panel of amphipathic peptides ranging from ApoA-I mimetic peptides to antimicrobial peptides (Fig. 1b, Extended Data Fig. 1a and Supplementary Table 1). Formation of MalFGK_2_ NDs from direct incubation of proteoliposomes with peptides was characterized using blue-native electrophoresis (native PAGE) (Fig. 1b and Extended Data Fig. 1a). Once extracted from lipid bilayers into micelles or incorporated into NDs, MalFGK_2_ migrated as a single membrane protein complex with a molecular weight of ∼200 kDa, as observed with the detergent DDM. This simple readout enabled us to rapidly screen more than a dozen amphipathic peptides for detergent-free reconstitution of MalFGK_2_ into NDs. The results showed that several ApoA-I mimetic peptides (e.g., DeFr_1-3, Fig. 1b) were able to transform MalFGK_2_ proteoliposomes into NDs as compared to other amphipathic or antibacterial peptides. However, these NDs contained a mixture of monomer, dimer, and different oligomers of MalFGK_2_ since multiple bands of protein complexes with higher molecular weights were observed on the native PAGE.

Because DeFr_1 is the most effective for the extraction of MalFGK_2_ monomer, we then attempted to purify MalFGK_2_ NDs formed by DeFr_1 through size-exclusion chromatography (SEC) (Fig. 1c). As expected, soluble particles were isolated after incubating MalFGK_2_ proteoliposomes with DeFr_1. In control experiments, we only observed large, insoluble proteoliposomes in the absence of peptides. Quantitative analysis also confirmed the presence of ∼100 copies of lipids per ND (Extended Data Fig. 1b), consistent with the idea that DeFr_1 directly transforms proteoliposomes into discoidal particles. Moreover, MalFGK_2_ encased in DeFr_1 NDs is functional as its ATPase activity remains coupled to maltose and MBP (Fig. 1d), in line with previous studies obtained using proteoliposomes^42^. In contrast, MalFGK_2_ was inactive and not responding to maltose and MBP after extraction into NDs using amphipathic polymers (Extended Data Fig. 1c and d). The lack of detectable ATPase activities of MalFGK_2_ in polymer NDs is probably due to the sensitivity of these polymers to bivalent cations^29^.

We further characterized the DeFr_1 enclosed MalFGK_2_ NDs using negative-stain electron microscopy (EM). Although most purified particles from SEC are indeed showing diameters of 10-20 nanometers (Fig. 1e), they are quite poly-disperse, consistent with the data from native PAGE analyses (Fig. 1b). In addition, the structural feature of the MalFGK_2_ transporter is difficult to identify from these raw micrographs, suggesting that DeFr_1 needs further engineering to improve its performance in ND reconstitution for biophysical studies of membrane proteins. For simplicity thereafter, we named these peptides DeFrMSPs for the reconstitution of <u>de</u>tergent-<u>fr</u>ee NDs through amphipathic <u>m</u>embrane <u>s</u>caffold <u>p</u>eptides.

### Potentiation of DeFrMSPs for ND reconstitution

Next, we sought to improve the efficacy of DeFr_1 for the reconstitution of native NDs. The key lies in the ability of the peptide to insert itself into lipid bilayers and stably enclose membranes, which can be potentiated by N- and C-terminal modifications (Extended Data Fig. 2a). First, we attempted to fuse DeFr_1 with antimicrobial peptides to enhance its membrane insertion activities. Second, we set out to functionalize DeFr_1 with distinct chemical groups, such as amide, acetyl, and fatty acids. To assay the efficiency of native ND reconstitution, the same MaFGK_2_ proteoliposomes were incubated with an increasing amount of these modified peptides and analyzed by native electrophoresis. The results showed that chemical modifications of DeFr_1 profoundly enhanced the monodispersity of MalFGK_2_ NDs (Fig. 2a and Extended Data Fig. 2a), migrating as a single band on non-denaturing native gels. Among all the different modifications, DeFr_5 resulted in the highest efficiency and was selected for further characterization.

**Figure 2.**
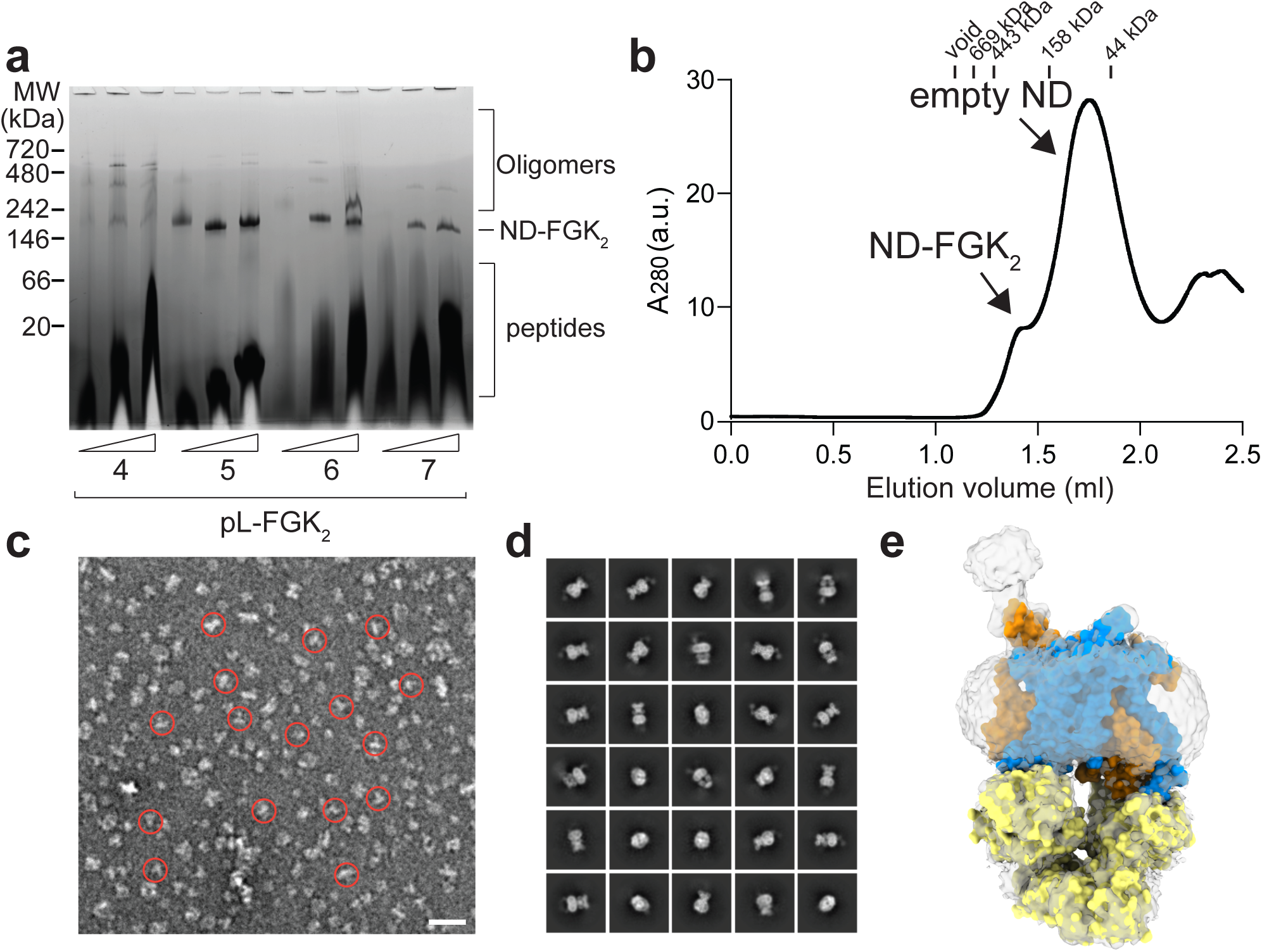
Chemical modifications enhance the efficacy of DeFrMSPs. (**a**) Blue native gel analysis of detergent-free ND reconstitution. MalFGK_2_ proteoliposomes prepared using PC lipids were incubated with the indicated peptide scaffolds (33, 100 and 300 µM) modified with fatty acids in the absence of detergents and then analyzed by native electrophoresis. (**b**) SEC profiles of detergent-free MalFGK_2_ NDs formed using of DeFr_5. Samples were fractionated on a Superdex S200 3.2/300 column. (**c**) Representative negative stain EM micrograph of MalFGK_2_ NDs encased by of DeFr_5. Scale bar, 30 nm. MalFGK_2_ NDs were highlighted in red circles. (**d-e**) Single particle cryoEM analysis of MalFGK_2_ in DeFr_5-enclosed NDs. d, 2D class average; e, 3D reconstructed model overlaid with density map (white).

We purified MalFGK_2_ NDs enclosed by DeFr_5 from SEC and determined their ATPase activities (Fig. 2b and Extended Data Fig. 2b). Again, the transporter is fully functional and stimulated by maltose and MBP. In addition, lipids were coeluted with these NDs (Extended Data Fig. 2c), as expected for the conversion of MalFGK_2_ proteoliposomes into ∼10-15 nm discoidal particles. We then analyzed these samples by negative stain EM (Fig. 2c). In contrast to the results obtained with DeFr_1, NDs formed by DeFr_5 were much more monodisperse, with clearly appreciable features of the cytosolic ATPase subunit of MalFGK_2_. The diameters of these MalFGK_2_ NDs formed with DeFr_5 averaged at ∼12 nm, which would provide a ∼2.5 nm layer of PC lipids around the transporter.

The vastly improved homogeneity of the DeFr_5 MalFGK_2_ NDs suggested that these samples are suitable for structural studies using single particle analysis by cryogenic electron microscopy (cryoEM). Indeed, the structure of the MalFGK_2_ complex in DeFr_5 NDs was solved in the closed state at 3.26 Å in the absence of ATP (Fig. 2d and e, Extended Data Fig. 3, and Supplementary Table 2), and is overall in agreement with previous crystal structures of the transporter stabilized with truncations in the N-terminus of MalF or arrested by the glucose enzyme EIIA^43, 44^. Together, we concluded that DeFr_5 can extract MalFGK_2_ into DeFrMSP-enclosed, detergent-free NDs (DeFrNDs) from membranes for biochemical and biophysical characterizations.

### DeFrNDs enclose stable and functional membranes

Although the above results obtained with MalFGK_2_ as a proof-of-principle study are encouraging, the properties of membranes in DeFrNDs remain unclear. In many cases, previous studies have shown that the stability and function of membrane mimetics vastly affect the conformational state of membrane proteins^45–47^. Furthermore, the lipids in polymer NDs rapidly diffuse from one ND to another by collisional transfer, unlike lipids in vesicles or traditional NDs formed by protein scaffolds^30^. This fast exchange of lipids between NDs implies that the polymer NDs may be rather dynamic and do not fully recapitulate the stable environment of cell membranes. Therefore, it is critical to assess the membrane property of DeFrNDs.

We employed a classic FRET-based membrane fusion assay to determine if the lipids in DeFrNDs are stably embedded or readily exchanging among individual NDs^48^. In these experiments (Fig. 3a), donor NDs or liposomes were prepared with a FRET pair (NBD-PE and Rho-PE) and then incubated with acceptor liposomes that did not harbor fluorescent lipids^49, 50^. If the lipids in DeFrNDs are unstable, the FRET pair, upon incubation with acceptor liposomes, will be diluted and separated from each other, increasing NBD fluorescence. However, we only observed a negligible fluorescence increase after incubating protein-free donor DeFrNDs with acceptor liposomes (Fig. 3b), similar to the negative control conditions using protein-free donor liposomes. In positive control experiments, we employed the classic membrane fusion machinery, soluble N-ethylmaleimide-sensitive-factor attachment protein receptors (SNAREs). Specifically, the cognate vesicle (v) and targeted-membrane (t) SNAREs were reconstituted into donor NDs and acceptor liposomes, respectively. Membrane fusion between NDs and liposomes mediated by the SNARE complex resulted in the massive dilution of the FRET pair and the dequenching of NBD fluorescence. Thus, lipids are stably enclosed in DeFrNDs for the characterization of membrane dynamics and do not freely diffuse between NDs unless fusogenic proteins are present.

**Figure 3.**
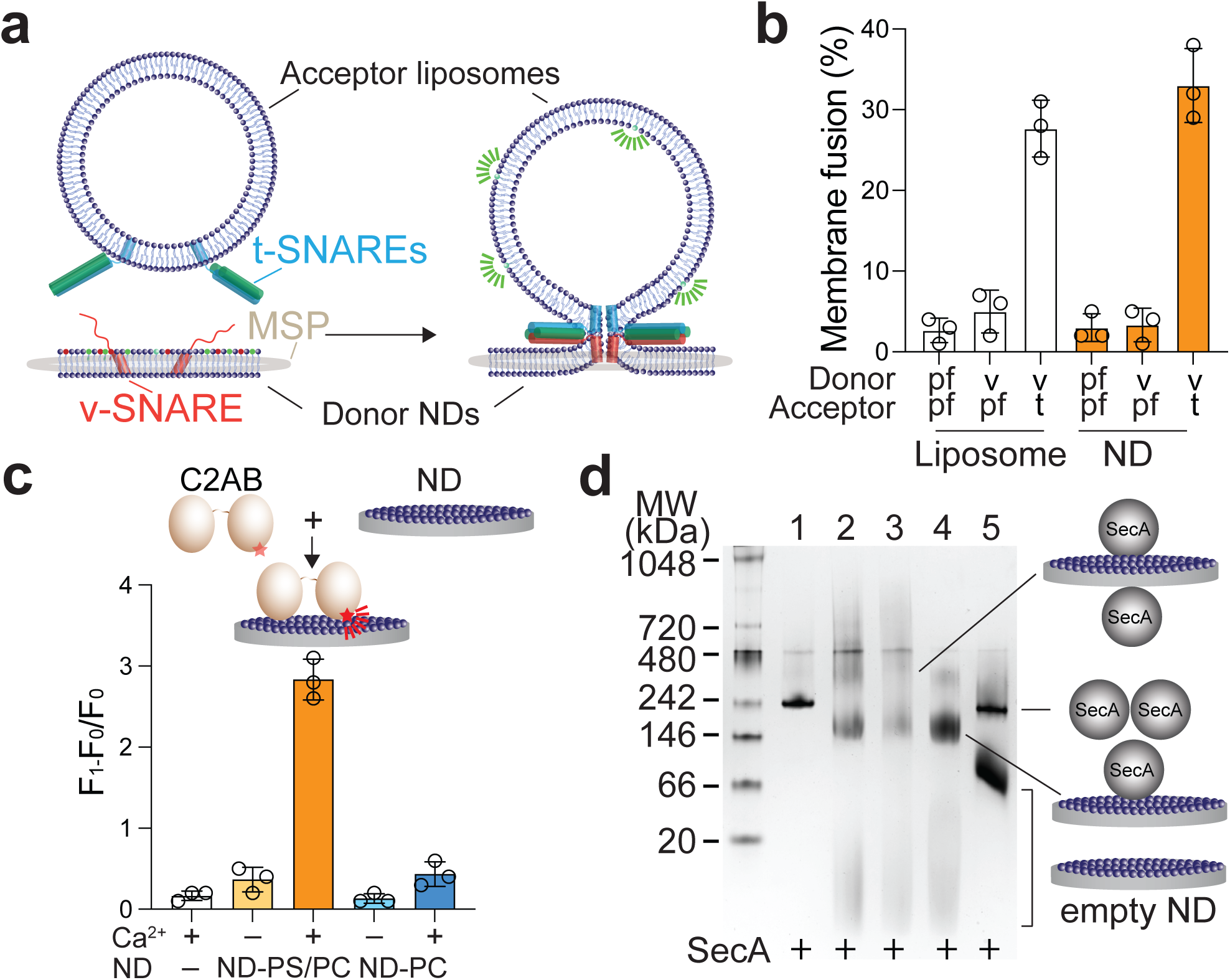
DeFrMSPs enclose stable and functional membranes. (**a**) Illustration of the membrane fusion assay. Donor NDs harboring NBD-PE and Rho-PE are incubated with acceptor liposomes. Fusion mediated by SNARE proteins will result in robust dequenching of NBD fluorescence. (**b**) Fusion activities of DeFrMSP NDs. Fusion assays were carried out using protein-free (pf) or v-SNARE (v) liposomes and NDs (donor, 0.5 µM) with liposomes (acceptor, 5 µM) that either contained t-SNARE (t) or were pf. Fusion assays were performed in the presence of Ca^2+^ and the soluble fragment of syt1. Data are shown as mean ± s.d., n = 3 independent experiments. (**c**) Characterization of syt1-lipid interactions using DeFrMSP NDs. Illustration (top) and quantification (bottom) of the syt1 membrane binding and penetration assay. The soluble fragment of syt1 (C2AB, 10 nM) was labeled with NBD and incubated with NDs (5 µM) encasing the indicated lipids (ND-PS/PC and ND-PC) in the presence or absence of Ca^2+^. Fluorescence of C2AB-NBD was quantified before (F_0_) and after (F_1_) the addition of NDs. Data are shown as mean ± s.d., n = 3 independent experiments. (**d**) Asymmetric membranes trapped in DeFrNDs as demonstrated by SecA-lipid interactions. SecA (1 µM) was incubated with DeFrNDs (2 µM) formed with the indicated lipid composition. Samples were then analyzed by clear native electrophoresis. 1, SecA alone; 2, SecA with NDs prepared with only PG lipids (ND-PG); 3, SecA with ND prepared with symmetric PG and PC lipids (sym ND-PG/PC); 4, SecA with NDs prepared with asymmetric PG and PC lipids (asym ND-PG/PC); 5, SecA with NDs prepared with only PC lipids (ND-PC).

Finally, we assessed if the DeFrNDs can functionally capture protein-lipid interactions. For this purpose, NDs were prepared with different lipid mixtures to characterize membrane binding and remodeling by synaptotagmin-1 (syt1), the Ca^2+^ sensor responsible for synaptic transmission^51–53^. Using a well-established membrane penetration assay^54^, syt1-lipid interactions will cause a drastic fluorescence increase only in the presence of Ca^2+^ and anionic lipids, as we found using DeFrNDs (Fig. 3c). Consistently, minor increases were observed in the absence of Ca^2+^ or with NDs harboring solely neutral lipids. Together, we conclude that DeFrNDs can encase stable and functional lipid bilayers for biophysical characterizations of membrane biology.

### Exploring DeFrNDs to trap membrane asymmetry

One distinct feature of native membranes is lipid asymmetry that is critical for numerous cellular signaling pathways^55, 56^. However, it is challenging to reconstitute this lipid asymmetry in traditional NDs because detergent-mediated reconstitution will disrupt the structural organization of membranes. We suspect that our peptide-mediated formation of native NDs might help address this dilemma by obviating the need for detergent solubilization.

We first formed asymmetric PC/PG liposomes using a previously described pH-driven method^57^ and incubated these liposomes with DeFr_5 to form NDs. If membrane asymmetry is preserved in DeFrNDs, they should contain mainly PG lipids on one side and largely PC lipids on the other side. We then used SecA, which has a high affinity for PG lipids, to assess the asymmetry of PG and PC lipids in DeFrNDs (Fig. 3d). SecA is a ∼100 kDa protein and forms a dimer of ∼200 kDa in solution. Interestingly, it dissociates into monomers upon binding to negatively charged lipids^58^. By incubating SecA with DeFrNDs harboring only PG or symmetric PG/PC lipids (Lane 2 and 3), we thus readily observed two additional bands showing up on native gel electrophoresis. The upper band was the complex of two SecA monomers bound to each side of NDs. The lower band corresponded to one monomeric SecA with NDs, which migrated below the SecA dimer band probably because of the negative charge of PG lipids. In contrast, SecA did not bind to PC lipids, so DeFrNDs prepared with only PC lipids (lane 5) had little impact on the migration of the protein on the native gel compared to SecA alone (lane 1). With asymmetric membranes (lane 4), the lower band of the ND-SecA monomer complex was much more enriched, indicating that membrane asymmetry was perhaps reconstituted in DeFrNDs.

### Reconstitution of bacterial membrane protein complexes into DeFrNDs

In contrast to detergent-mediated reconstitution, the main advantage of DeFrNDs is the potential to stabilize membrane protein complexes with native lipids (Fig. 4a). To test this idea, we incubated crude membranes from *E.coli* expressing MalFGK_2_ with DeFr_5 and performed affinity purification. As expected, we can directly isolate MalFGK_2_ in DeFr_5 -encased native NDs (Fig. 4b and Extended Data Fig. 4a). However, we noticed that DeFr_5 was much less efficient than the detergent DDM to extract MalFGK_2_, albeit significantly higher than DeFr_1 or negative control experiments (Fig. 4b and Extended Data Fig. 4a). To further enhance the performance of DeFr_5, we set out to optimize the peptide sequence. The amphipathic peptide DeFr_5 was designed to combine positively charged residues with hydrophobic ones. The main membrane penetrating residue of DeFr_5 is Phe. However, it is known that Trp is more potent to destabilize lipid bilayers. Therefore, we gradually replaced Phe with Trp and assessed their efficacies in extracting MalFGK_2_ into NDs from proteoliposomes. These peptides are all chemically modified as DeFr_5. In addition, we slightly increased the amphipathic repeat of DeFr_5 to 20 or 22 amino acids to expand the surface area of these peptides for engaging lipids. As compared to DeFr_5, several of these redesigned peptides showed improvement in extracting MalFGK_2_ into NDs from proteoliposomes (Extended Data Fig. 4b), with DeFr_12 exhibiting the highest efficacy. We then placed Trp substitutions in DeFr_12 and assessed their performance for the reconstitution of MalFGK_2_ NDs from crude *E.coli* membranes. Consistent with the results obtained using proteoliposomes, Trp substituted DeFr_14 peptides were indeed more effective than the original DeFr_5, giving rise to a ∼2.5-fold increase in the yield of native NDs (Fig. 4b). Because DeFr_14 showed the highest reconstitution efficiency, we purified native NDs formed with this peptide for further characterizations. These MalFGK_2_ NDs encased by DeFr_14 were quite homogenous as shown by SEC, negative stain EM and dynamic light scattering (DLS) analyses (Fig. 4c and Extended Data Fig. 4c-e) and contained native bacterial lipids (Extended Data Fig. 4f). The associated lipids with MalFGK_2_ were mainly PE and PG. The absence of other bacterial lipids, such as cardiolipin, indicated that the transporter was surrounded by a specific local lipid environment.

**Figure 4.**
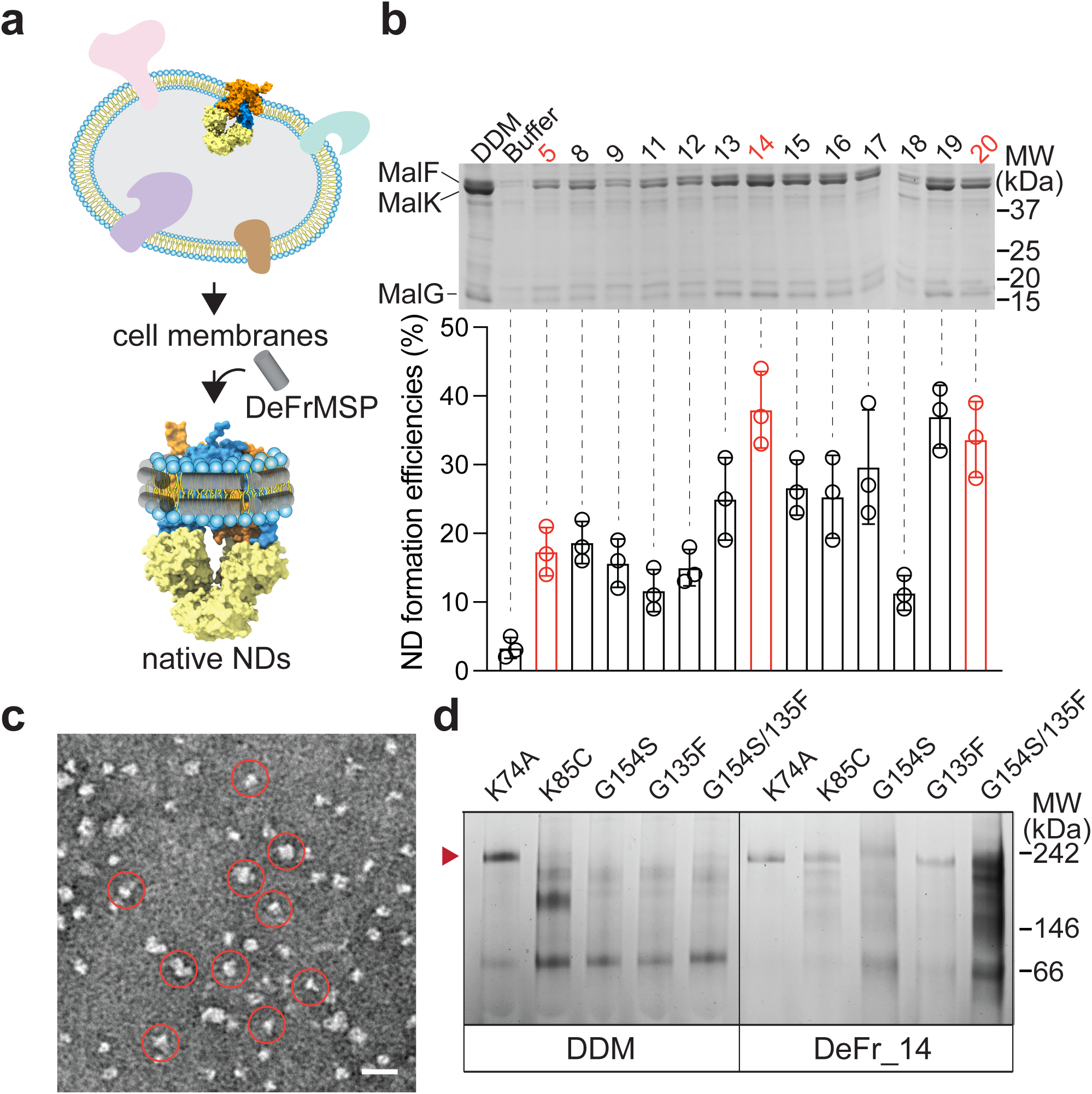
Formation of native NDs from bacterial membranes using DeFrMSPs. (**a**) Illustration of the detergent-free, native ND reconstitution procedure. Native NDs were formed by one-step incubation of crude cell membranes with the designed DeFrMSPs in the absence of detergents. (**b**) ND formation efficiencies of MalFGK_2_ with different designs of DeFrMSPs. DDM and PBS buffer were used as positive and negative controls for data normalization. Top: representative gels of extracted MalFGK_2_ in native NDs. Bottom, ND formation efficiencies of MalFGK_2_ were quantified based on the density of the MalF and MalK bands normalized to that of DDM-extracted samples. Data are shown as mean ± s.d., n = 3 independent experiments. (**c**) Negative stain EM micrograph of MalFGK_2_ NDs encased by DeFr_14. Native MalFGK_2_ NDs were highlighted in red circles. Scale bar, 30 nm. (**d**) Blue native gel analysis of isolated MalFGK_2_ mutants in DDM or DeFr_14 NDs. Red arrow indicates the position of the intact MalFGK_2_ complex. Most of these mutant complexes were unstable once extracted by DDM from cell membranes and thus dissociated into individual subunits. In contrast, many of them can be stably extracted into native NDs by DeFr_14.

Since our DeFrNDs can retain native lipids, they might also be able to trap challenging membrane protein complexes that are otherwise disrupted by detergents. To test this idea, we assayed the performance of DeFr_14 to isolate MalFGK_2_ mutants that fall apart once extracted out of membranes by detergents^60^. These mutants behave very differently from the wild-type protein and are useful tools to advance our understanding of ABC transporters in general^61^. However, in-depth dissections of these mutants are unattainable because they are not stable in detergents. As shown by native electrophoresis, several MalFGK_2_ mutants were dissociated at various degrees after affinity purification in DDM (Fig. 4d, left). In contrast, DeFr_14 directly extracted many of these MalFGK_2_ mutants into NDs with high efficiency and good stability for biochemical characterizations (Fig. 4d, right). We found that the basal ATPase activities of these mutants were much higher than the wild-type MalFGK_2_ transporter (Extended Data Fig. 4g), indicating escalated alternations in their conformational dynamics for MBP-independent maltose translocation.

### Reconstitution of eukaryotic membrane proteins into DeFrNDs

To further assess the utility of DeFrNDs for biophysical characterizations of membrane biology, we first explored if they are appropriate tools for reconstituting a human GPCR, the metabotropic glutamate receptor 7 (mGluR7). We expressed the receptor from HEK293 cells and assayed if we could extract it from crude cell membranes into native NDs using DeFrMSPs (Fig. 5a and Extended Data Fig. 5a). We genetically fused mGluR7 with eGFP, allowing for sensitive fluorescent measurements to determine the efficiency of ND formation. We then incubated our collection of DeFrMSPs with mGluR7-eGFP membranes and quantified the efficiency of NDs formation by measuring the fluorescence of eGFP in the solubilized fractions (Extended Data Fig. 5a). The formed native NDs were then purified and analyzed using denaturing and native electrophoresis followed by in-gel fluorescence imaging (Fig. 5b and Extended Data Fig. 5b). We found that DeFr_14 was the most effective to extract mGluR7 into soluble and homogeneous NDs as a single band of the reconstituted complex was observed on native PAGE (Fig. 5b). We purified mGluR7 native NDs formed by DeFr_14 through affinity purification and SEC (Extended Data Fig. 5b and c). Using negative stain EM and DLS measurements, we found that mGluR7 native NDs are indeed quite monodisperse. Moreover, they are fully functional to bind with high affinity to the extracellular domain of the trans-synaptic binding partners of mGluR7^62^, the extracellular leucine-rich repeat and fibronectin type III domain-containing protein (ELFN-1), as shown by native electrophoresis (Fig. 5b and c).

**Figure 5.**
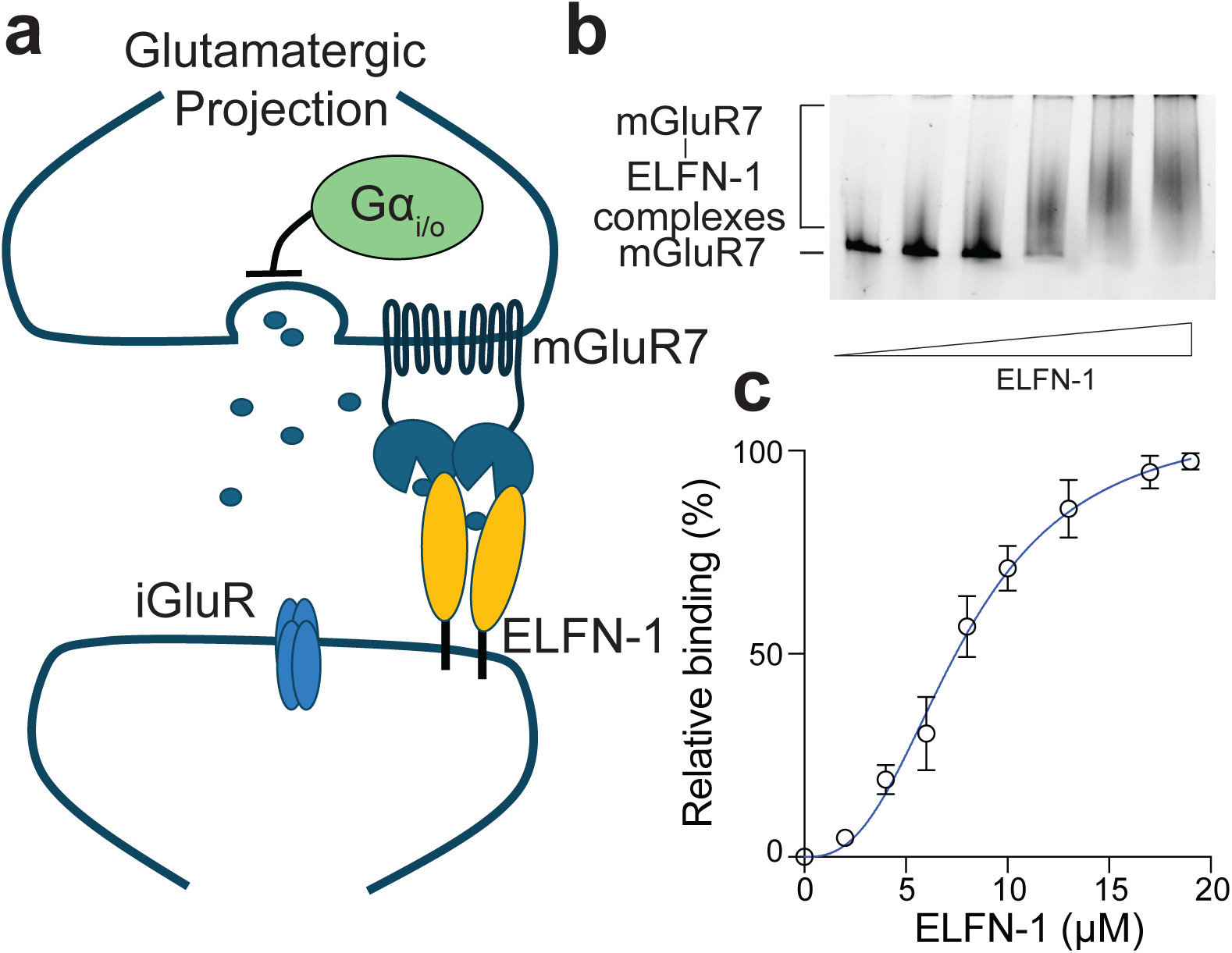
Reconstitution and characterization of mGluR7 in native NDs. (**a**) Illustration of the mGluR7-ELFN complex at neuronal synapses. mGluR7 is a prominent metabotropic glutamate receptor that modulates synaptic transmission. Recent studies demonstrate that its function is also regulated by trans-synaptic interactions with ELFN-1^62^. (**b**) Representative clear native gel of mGluR7 in DeFr_14 NDs binding to ELFN-1. Native NDs harboring mGLuR7 were incubated with increasing concentrations of the extracellular fragment of ELFN-1. Samples were analyzed by native electrophoresis and in-gel fluorescence imaging. (**c**) Quantification of mGluR7 interaction with ELFN-1 from native gel analysis as described in b. The decreased densities of the mGluR7 band normalized to the protein alone (lane 1) was used to calculate the relative binding to ELFN-1 at various concentrations. The binding data of mGluR7 with ELFN-1 was then fitted with the Hill equation (*K*_d_ = 8 ± 1 µM; n= 2.6). Data are shown as mean ± s.d., n = 3 independent experiments.

In addition to mGluR7, we also tested the performance of our engineered DeFrMSPs for native ND reconstitution of the HCN1 channel (Fig. 6). Consistently, we can extract HCN1 into NDs from crude HEK293 membranes (Extended Data Fig. 6a), as characterized by SEC, negative stain EM and DLS (Extended Data Figs. 6b and c). Interestingly, DeFr_5 was the best scaffold for the extraction of HCN1, indicating that its local lipid environment might be different from mGluR7. The SEC profile of HCN native NDs showed more oligomerized proteins than mGluR7, perhaps because the fused GFP on this tetrameric channel tends to self-associate into high-order oligomers in the absence of detergents. Using a powerful single-molecule fluorescence imaging assay in zero mode waveguides^63^, we looked at the binding activity of fcAMP to HCN1 channels (Fig. 6a and b). The results showed that the HCN1 tetramer is functional in DeFrMSP-encased native NDs, retaining the ability to bind fcAMP (Fig. 6c and d). The overall binding activity and affinity of HCN1-cAMP interactions in native NDs were very different from data obtained in detergent micelles, supporting the role of lipids in regulating the property of this ion channel. In contrast, we could not efficiently purify HCN1 native NDs using amphipathic polymers (Extended Data Fig. 6d), let alone maintain their activities. We noted that the presented model in Figure 6b is a simplified illustration of cAMP binding to HCN1. The underlying molecular mechanism is a subject of debate and will require more extensive characterization to generate in-depth insights into the interaction of cAMP with HCN1.

**Figure 6.**
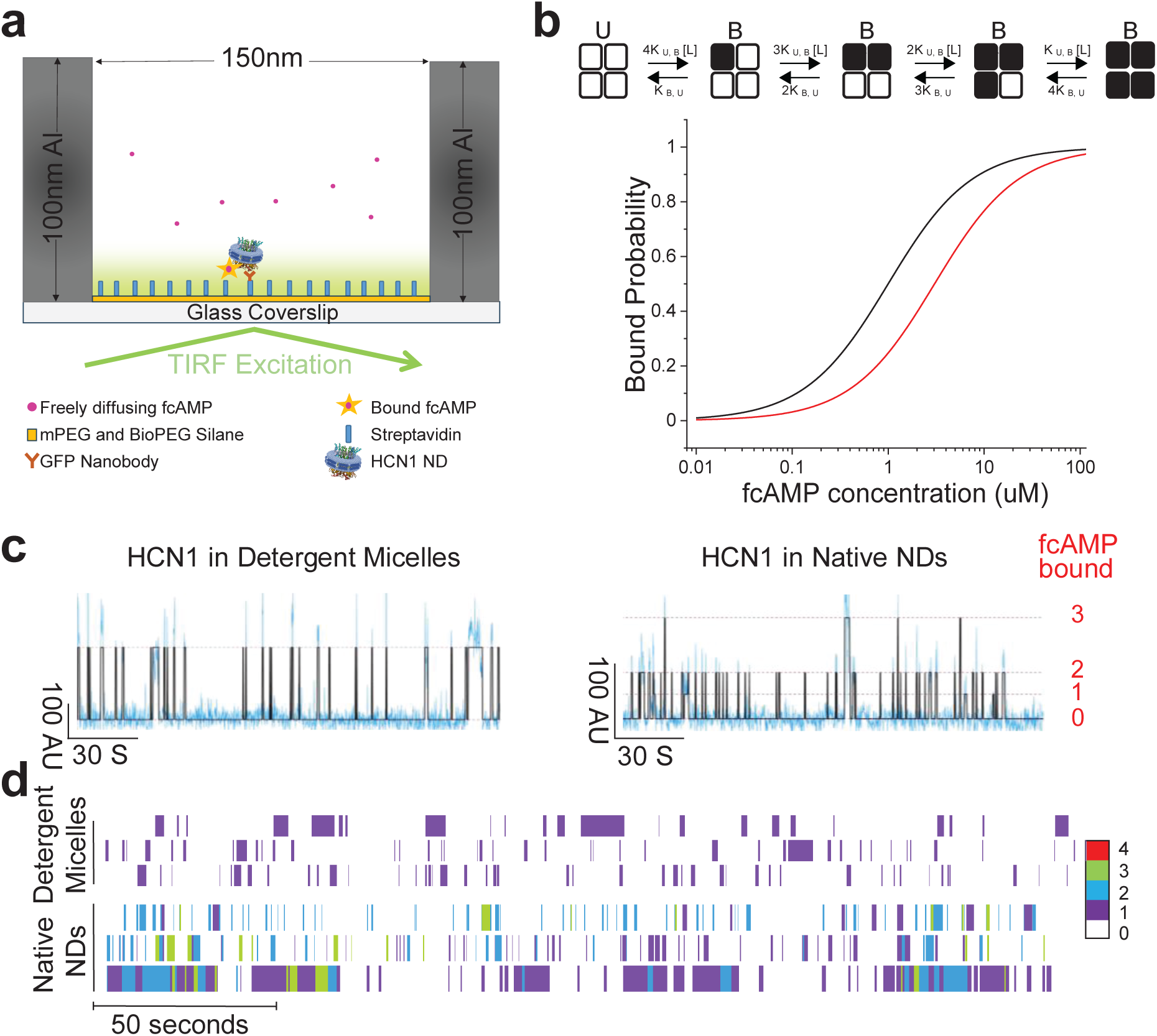
Reconstitution and characterization of HCN1 in native NDs. (**a**) Illustration of the single molecule fluorescence ligand binding assay to probe the interaction of fcAMP with HCN1. (**b**) fcAMP binds to HCN1 in native NDs with higher affinity than HCN1 in detergent micelles. Top, independent sequential ligand binding model for fcAMP binding, white boxes indicate the unbound state (U) and black boxes (B) indicate the bound state. “L” represents the ligand and “*K*” represents the transition rate constant. Bottom, predicted binding curves for HCN1 in detergent (red) and HCN1 in native NDs (black) were generated by calculating the bound probability at various concentrations by using the Adair equation (see Methods). The binding probability of the first ligand was obtained from experimental data, and the binding probabilities of the 2^nd^, 3^rd,^ and 4^th^ ligands were estimated, assuming that these binding events occur independently. (**c**) Representative fluorescence-time traces overlaid with idealized fit showing fcAMP (50nM) binding to HCN1 in detergent micelles (left) or native ND (right). (**d**) Representative fluorescence-time traces displayed as heat maps show multiple traces of fcAMP (50nM) binding to HCN1 in detergent micelles (upper) or native ND (bottom). White spaces indicate unbound state of the channel whereas the color coding indicates duration and transition to various ligation states - 1 bound 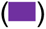, 2 bound 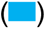, 3 bound 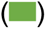 or 4 bound 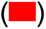. Across all concentrations, for detergent sample, total N= 386 molecules, total time – 35 hours, total events-35661. For HCN1 in native NDs, total N= 119 molecules, total time -9.9 hours, total events -13143.

Moreover, we performed thin layer chromatography (TLC) experiments to analyze the extracted lipids in native NDs (Extended Data Fig. 6e and f). Interestingly, we found that mGluR7 was associated with more PE lipids than HCN1. This difference might suggest the role of lipids in regulating the function of membrane proteins. Further, these results demonstrate that DeFrMSP NDs can isolate surrounding native lipids with eukaryotic membrane proteins.

## Discussion

In this manuscript, we engineered membrane scaffold peptides, DeFrMSPs, for detergent-free reconstitution of membrane proteins into NDs from native membranes. This DeFrND approach bypasses the limitation of detergent-mediated reconstitution used in traditional methods (Extended Data Fig. 7), thereby enabling structural and functional studies of membrane protein-lipid complexes previously unattainable. In addition, the membranes in DeFrNDs can more closely mimic native lipid environments for biophysical dissections of both integral and periphery membrane proteins involved in various transmembrane signaling pathways. We anticipate that this new approach will nicely complement current ND platforms and serve as a useful tool for both basic and translational research of membrane proteins.

While previous studies on ApoA1 mimetic peptides have established that some of them can solubilize synthetic membranes with defined lipid compositions^32, 33, 37, 64^, our engineered peptides can extract membrane protein-bearing discoids with high efficiency from native membranes. The reconstituted signaling membrane proteins (channels, receptors, and transporters) are never exposed to detergents throughout the reconstitution process, and the resultant NDs maintain their structure, function and single molecule dynamics (Figs. 2-6). In addition, DeFrMSPs are compatible with various membrane proteins and vastly simplify the workflow for the reconstitution of NDs compared to previous methods. The extraction activity of DeFrMSPs mostly depends on two parameters: insertion into membranes and stabilization of discoidal ND structures. Specifically, these peptides first need to penetrate lipid bilayers and then stably encircle a rigid patch of membranes into the ND framework. Antimicrobial peptides can readily insert themselves into membranes but cannot form stable discoidal structures with lipids. Hence, they are unable to extract membrane proteins into NDs as compared to ApoA-I mimetic peptides such as DeFr-1. In contrast, longer amphipathic peptides (e.g., NSP and NSPr) can form stable NDs^34^. However, they are much less effective in penetrating lipid bilayers to allow for detergent-free ND reconstitution (Fig. 2a, Extended Data Figs. 1a and 2a), probably due to the increased sizes. Consistently, it is known that traditional membrane scaffold proteins derived from ApoA-I are completely unable to disrupt membranes, most likely because of their over 10-fold larger molecular weights. Therefore, the successful development of DeFrMSPs is realized by optimizing membrane insertion and ND stability through extensive screening and rational design.

Further, DeFrMSPs greatly complement native ND reconstitution systems. Current native NDs are formed using different kinds of polymers that often require careful optimization^7, 27^, such as the concentration of divalent ions and pH in the reconstitution buffer. In addition, it is difficult to predict the compatibility of these polymers for a specific membrane protein, because their efficiencies for forming native NDs vary significantly from case to case. The compatibility of DeFrNDs with bivalent cations makes it a great complimentary system to current detergent-free reconstitution systems based on polymers. For example, DeFrNDs are suitable for membrane proteins that are not active in polymer-based NDs (Figs. 1-3 and Extended Data Fig. 1). However, we also noted that the current polymer-based system provides more options and versatility than DeFrND. Despite rapid lipid exchange between polymer NDs, they are stable and can withstand multiple free-thaw cycles^65^.

Moreover, DeFrMSPs will extend the potential of NDs for fundamental research of membrane biology. From our assessment of DeFrNDs with different transporters, receptors and ion channels, we found that the size of DeFrNDs adapted to the embedded transmembrane domain. Another advantage of DeFrND is the potential to reconstitute asymmetric membranes with synthetic and native lipids. By virtue of this advantage, further improvement of our approach might enable a variety of biophysical approaches to investigate how lipid asymmetry regulates the conformational dynamics of membrane protein complexes as observed in cells. In conjunction with single particle cryoEM, we expect that the DeFrND technology will help to unravel the molecular mechanism underlying the regulation of myriad transmembrane signaling pathways by native cell membranes. Based on our cryoEM analysis, DeFrNDs have not exhibited preferred orientation as sometimes found in other membrane mimetic systems^66^. This advantage will provide a useful alternative option for structural characterizations of membrane proteins in a near native lipid environment using single particle cryoEM.

DeFrMSPs will also benefit therapeutic developments based on ApoA-I mimetic peptides. NDs formed with ApoA-I mimetic peptides are potent vehicles for vaccination^14, 16–18^, because of their nanoscale sizes, enhanced tissue penetration, and low immunogenicity. As such, these nanomaterials can avidly engage with antigen-presenting cells and reprogram the immune system to control cancer and viral infections. We expect that the enhanced stability and efficiency of detergent-free ND reconstitution by DeFrMSPs will further promote therapeutic applications in these areas. Another potential application is to facilitate the delivery of therapeutic protein complexes (e.g., genome-editing enzymes)^67, 68^. In conjunction with cationic cell-penetrating peptides, several amphipathic peptides were recently engineered to bind protein therapeutics and boost their entry into cells through the unconventional clathrin-independent endocytosis and macropinocytosis. The delivery efficiency is determined by the membrane penetrating activities of these amphipathic peptides. Since we found that DeFrMSPs also could spontaneously insert themselves into cell membranes, we expect that fine-tuning of our peptide designs will generate robust platforms for delivering therapeutic proteins.

Together, our study establishes a new approach for native ND reconstitution through the development of membrane scaffold peptides, DeFrMSPs. The application of this powerful approach will elucidate how the structural organization of native membranes governs the dynamics and function of membrane protein complexes, thereby shedding light on new ideas to steer membrane biology for therapeutic developments against human diseases. Further improvement and optimization are critical to expand the utility of DeFrMSPs. One of the main challenges is the lower extraction efficiency of DeFrMSPs than detergents. This problem will need refinement of DeFrMSP designs to increase their solubilities and efficacies in extracting membrane proteins from native biological membranes. More design strategies can be learned from the characterization of novel amphipathic peptides and proteins^69^. Furthermore, we expect that machine learning and computational protein design approaches will greatly help our efforts in improving the performance of DeFrMSPs, as evidenced by the discovery of new potent antimicrobial peptides^70^.

## Methods

### Chemicals and Reagents

1,2-dioleoyl-sn-glycero-3-phosphoethanolamine-N-(7-nitro-2-1,3-benzoxadiazol-4-yl) (NBD-PE), 1,2-dioleoyl-sn-glycero-3-phosphoethanolamine-N-(lissamine rhodamine B sulfonyl (Rho-PE), 1,2-dioleoyl-sn-glycero-3-phosphocholine (PC), 1,2-dioleoyl-sn-glycero-3-phospho-l-serine (PS), 1,2-dioleoyl-sn-glycero-3-phosphoethanolamine (PE) and 1,2-dioleoyl-sn-glycero-3-phospho-(1’-rac-glycerol) (PG) were obtained from Avanti Polar Lipids. Nitrilotriacetic acid (Ni^2+^-NTA)-chelating Sepharose, Superdex 200 3.2/300 GL, Superdex 200 10/300 GL and Superose 6 increase 10/300 GL were purchased from GE Healthcare. 1,1’-Dioctadecyl-3,3,3’,3’-Tetramethylindocarbocyanine Perchlorate) (Dil), N,N’-Dimethyl-N-(Iodoacetyl)-N’-(7-Nitrobenz-2-Oxa-1,3-Diazol-4-yl)Ethylenediamine (IANBD amide) and Oregon Green^TM^ 488 (OG) maleimide were obtained from ThermoFisher. All other chemicals were acquired from Sigma.

### Plasmids

pTrc-MalFGK_2_ was a gift from Dr. Franck Duong^60^. pEThis-vamp2 was a gift from Dr. James Rothman^48^. pGEX-syt1 and pDuetRSF t-dimer were a gift from Dr. Edwin Chapman^50, 54^. All other constructs in this work were made using the In-Fusion® HD Cloning Kit (Takara Bio USA).

### Proteins

Protein expression and purification for syt1, MBP, SecA, MalFGK_2_ and SNAREs were performed as described previously^41, 49, 50, 71–73^. Briefly, plasmids were transformed into BL21 cells that were grown in LB supplemented with Km (50 mg/ml) or Amp (100 mg/ml) to OD_600_ ∼0.7. Protein expression was induced with 0.2 mM IPTG at 16 °C, overnight (O/N). Bacteria were harvested by centrifugation at 3, 428 x g for 20 mins, resuspended in Buffer A (50 mM Tris-HCl (pH 8),100 mM NaCl, 5% glycerol, 2 mM β-mercaptoethanol), and lysed using a Branson cell disrupter. Cell lysates were clarified by centrifugation at 8, 184 x g for 45 mins. For syt1, MBP and SecA, the supernatants were loaded onto a 1 ml NTA column (GE Healthcare), followed by two times wash using buffer B (50 mM Tris-HCl (pH 8), 20 mM Imidazole, 400 mM NaCl, 5% glycerol, 2 mM β-mercaptoethanol). Proteins were eluted in buffer C (50 mM Tris-HCl (pH 8), 500 mM Imidazole, 400 mM NaCl, 5% glycerol, 2 mM β-mercaptoethanol), desalted in buffer A using PD MiDiTrap G-25 (GE Healthcare) for syt1 and MBP, and stored at -80 °C. SecA was further purified by SEC using a Superdex 200 10/300 column in 50 mM Tris-HCl (pH 8), 100 mM NaCl, 5% glycerol, 1mM DTT. Soluble SecA dimer fractions were pooled together and concentrated to ∼2 mg/ml and stored at -80 °C. For MalFGK_2_ and SNAREs, membrane fractions were resuspended in Buffer A and solubilized with 1% DDM (4 °C, O/N). Solubilized membrane extracts were isolated by centrifugation at 112, 000 × g for 1 h. Supernatants were loaded onto a 1 ml NTA column (GE Healthcare), followed by two times wash using buffer B supplemented with 0.02% DDM. Proteins were eluted in buffer C plus 0.02% DDM, desalted in buffer A supplemented with 0.02% DDM using PD MiDiTrap G-25 (GE Healthcare) and stored at -80 °C.

Ecto-ELFN1-Fc (residues 1-399) were expressed as previously described^62^. An eGFP fusion was added to the C-terminus of mGluR7 and expressed as before^62^. Briefly, mGluR7 and Ecto-ELFN1-Fc constructs were inserted into modified pEG BacMam vector^74^. The bacmid was then transfected into Sf9 cells (ThermoFisher Scientific, 11496015) to produce baculovirus. The supernatant of the third generation of virus production (P3) was then harvested and added to HEK293S GnTi-cells (ATCC, CRL-3022) grown at 37 °C with 8% CO2 at a confluency of 2.0-3.0×10^6^ cells/mL in FreeStyle 293 expression media. After 18 hours, sodium butyrate (Sigma-Aldrich, B5887) was added to 10 mM. The cells were then incubated for a further 48 hours and pelleted by centrifugation. For the purification of ELFN-1, the supernatant of cell lysates clarified by centrifugation (8, 184 x g, 1 h) was then affinity purified using Pierce™ anti-DYKDDDDK affinity resin (ThermoFisher Scientific, A36801) and eluted with 3 x Flag peptide (GenScript, RP21087), concentrated using an Amicon™ Ultra-4 centrifugal filter (Sigma-Aldrich, UFC805024), analyzed by SDS-PAGE, aliquoted and flash frozen.

Expression of HCN1 tagged with eGFP and Single molecule fluorescence imaging were performed as described previously^63^.

### Fluorescent labeling of proteins

Purified syt1 were desalted using Zeba Spin columns (Thermo Fisher) in buffer D (50 mM Tris-HCl, pH 8, 100 mM NaCl, 5% glycerol) and labeled with a 3-fold excess of IANBD amide or OG maleimide in the presence of TCEP (0.2 mM) at room temp for 2h. Free dyes were removed by passing through Zeba Spin columns in buffer A.

### Negative stain electron microscopy

Formvar/carbon-coated copper grids (01754-F, Ted Pella, Inc.) were glow discharged (15 mA, 25 secs) using PELCO easiGlow^TM^ (Ted Pella, Inc). NDs (10 µg/ml) were applied onto the grids for 30 secs, followed by staining with 0.75% uranyl formate for 1 minute. Images were collected using a ThermoFisher Science Tecnai G2 TEM (100 kV) equipped with a Veleta CCD camera (Olympus). All TEM data were analyzed using Fiji to determine ND sizes.

### CryoEM data collection

3µl of MalFGK_2_ NDs at a concentration of 2mg/ml was applied to glow discharged 300 mesh UltraAufoil gold R1.2/1.3 (quantifoil) and vitrified using a Vitrobot Mark IV (Thermo Fisher Scientific/FEI) at 10 °C and 100% humidity with a blot time of 4s, and a blot force of 0. Cryo-EM data for single particle analysis were collected at New York Structural Biology Center on a 300 kV Titan krios electron microscope (Thermo Fisher Scientific/FEI) with a Gatan K3-Bioquantum direct electron detector (Gatan, Inc.) and energy filter, using leginon software^75^. Movies were captured at a magnification of 81,000x (pixel size of 1.058 Å). A total of 12522 movies in the defocus range of -0.8 to -2.2 μM were recorded with a total accumulated dose of 55.80 e-/Å.

### Image processing, 3D reconstruction, modeling, and refinement

Data processing was performed using cryoSPARC v4.4.1^76^. Raw movies were aligned using patch motion correction, and the micrograph contrast transfer function (CTF) parameters were estimated via patch CTF estimation. Micrographs were picked using a blob picker, and an initial particle set was chosen through 2D classification. Selected 2D classes were used for templet particle picking. Ab-initio jobs were run with 4 classes, and one of the best classes was used for further data processing. Three rounds of heterogeneous refinement were employed to remove junk particles. Details of the Cryo-EM classification and processing are shown in Extended Data Fig. 3. The resulting curated particle sets were used for local CTF refined to high resolution using non-uniform refinement with 3.26 Å resolution using the 0.143 Fourier Shell Correlation criterion.

A model was built starting from PDB structure of truncated 7t4e. The initio model was rigid body docked into the density and adjusted in Chimera^77^ and coot^78^. Loops were manually fit into the density map using coot. Subsequently, the real space refinement was performed, with remaining manual adjustments were performed in Coot. Models were validated using Molprobity in Phenix^79^. A summary of model refinement statistics is provided in Supplementary Table 2.

### Fusion assays

v-SNARE vesicles harboring the FRET reporter (NBD-/Rho-PE lipids) and t-SNARE vesicles were prepared as described previously^49, 50^. Fusion assays were performed by incubation of v-SNARE NDs or vesicles (0.5 µM) with t-SNARE vesicles (5 µM) in reconstitution buffer (20 mM Tris-HCl, pH 7.5, 100 mM NaCl) at 37 °C for 30 mins. For lipid exchange assays, NDs (0.5 µM) were prepared from protein-free liposomes bearing the same FRET reporter and incubated with another set of NDs (5 µM) prepared from protein-free liposomes using PC/PE/PS lipids as described above. NBD fluorescence was monitored using a Synergy H1M plate reader with excitation at 460 nm and emission at 530 nm. After each run, the percentages of membrane fusion or lipid exchange were calculated by normalization of data to the maximal fluorescence signal increase after the addition of 0.5 % DDM to each reaction.

### Fluorescence spectroscopy

The NBD-labeled soluble fragment of syt1 (C2AB, 10 nM) was incubated with the indicated NDs (5 µM). The fluorescence spectrum of samples was collected on a Synergy H1M plate reader with excitation at 460 nm and emission from 500 to 650 nm.

### Single molecule cAMP binding measurements

Single molecule cAMP binding measurements were done using zero-mode waveguides fabricated as described previously^63^. Imaging was done on a custom micromirror TIRF setup (Mad city labs) equipped with a NA 60X oil immersion objective lens (Olympus). Micromirror TIRF excitation field generated either with a 488nm or 561nm lasers (OBIS, coherent) to excite the GFP and fcAMP, respectively. The subsequent fluorescence emission was recorded on a 512×512 EMCCD camera (Andor iXON Ultra) at a frame rate of 10Hz. Data collection was done using Micromanager 2.0. The emission was passed filtered through a dichroic filter (T565lpxr Chroma) and then a band-pass filter (Chroma ET550/25 nm) for GFP and a band-pass filter (Semrock bright line 593/40nm) for fcAMP were in the corresponding emission pathways. The binding data were recorded from each molecule for a minimum of 240 seconds with 100 msec exposure time. Single-molecule data was analyzed as described in the previous work^63^. Briefly, single molecule traces were extracted and analyzed using the DISC software for the colocalization and image projection to obtain time-dependent fluorescence intensity changes. The traces were then idealized using the DISC software^80^. The equilibrium constant for intrinsic binding affinity, K, was calculated from the dwell times corresponding to the singly-bound state as described previously using QUB software^63^. Assuming that the binding of each of the four ligands binds to the channel independently, the binding curve was generated by calculating microscopic rate constants k1=K/4, k2=2/3(K), k3=3/2(K), and k4= 4K and substituting them in Adair’s equation^81^.

## Acknowledgment

Mass Spectrometry analyses were performed by the Mass Spectrometry Technology Access Center at the McDonnell Genome Institute (MTAC@MGI) at Washington University School of Medicine, supported by the Diabetes Research Center/NIH grant P30 DK020579, Institute of Clinical and Translational Sciences/NCATS CTSA award UL1 TR002345, and Siteman Cancer Center/NCI CCSG grant P30 CA091842. We thank Drs. Michael Purge and David Cooper at the University of Virginia and Drs. Naomi Kamasawa and Debby Guerrero-Given from the imaging center at the Max Planck Florida Institute for Neuroscience for assist in negative stain EM. CryoEM sample preparation and data collection were performed with the generous support from NCCAT. We thank Dr. Susovan Roy Chowdhury at WUSTL and Dr. Scott T. Retterer from Oak Ridge National Laboratory for generating ZMWs used in this study. We would also like to thank Dr. Franck Duong for the plasmid pTRC-MalFGK_2_, Dr. James Rothman for the plasmid pEThis-Vamp2, and Dr. Edwin Chapman for the plasmid pGEX4T-syt1-C2AB and pDuetRSF t-SNARE dimer. This work was supported by NIH (DP2GM140920 and R21AG078699 to H.B.; EY034339 to K.M.; NS124758 to W.G.L.; R35NS116850 to B.C.; P01GM072694 to L.K.T).

## Author Contribution

H.B. conceived the project and wrote the manuscript with input from all authors. Q.R., S.Z., and H.B. performed protein purification, biochemical reconstitutions, and characterizations of NDs by gel electrophoresis, SEC and ATPase assays. J.W., Q.R. and H.B. performed single particle analysis of NDs by cryoEM. J.S. and Q.R. performed negative stain EM analysis of NDs. H.C. contributed to the purification of SecA. V.I., S.R.C. and B.C. provided cells expressing HCN1 and carried out single molecule imaging experiments and data analysis, and lipidomic experiment and analysis. A.K., V.K. and L.K.T. provided dense core vesicles. S.Z. and A.J. performed extraction of Sec61β from ER microsomes. W.G.L. and K.M. purified ELFN-1 proteins and provided cells expressing mGluR7. L.H. and I. L. contributed to TLC experiments.

## Data availability

The coordinate and cryo-EM map of MalFGK_2_ NDs have been deposited at the PDB and Electron Microscopy Data Bank under accession code 9BCR (EMD-44435). All data supporting the findings of this study are available within this manuscript.

---

**Extended Data Figure 1.**
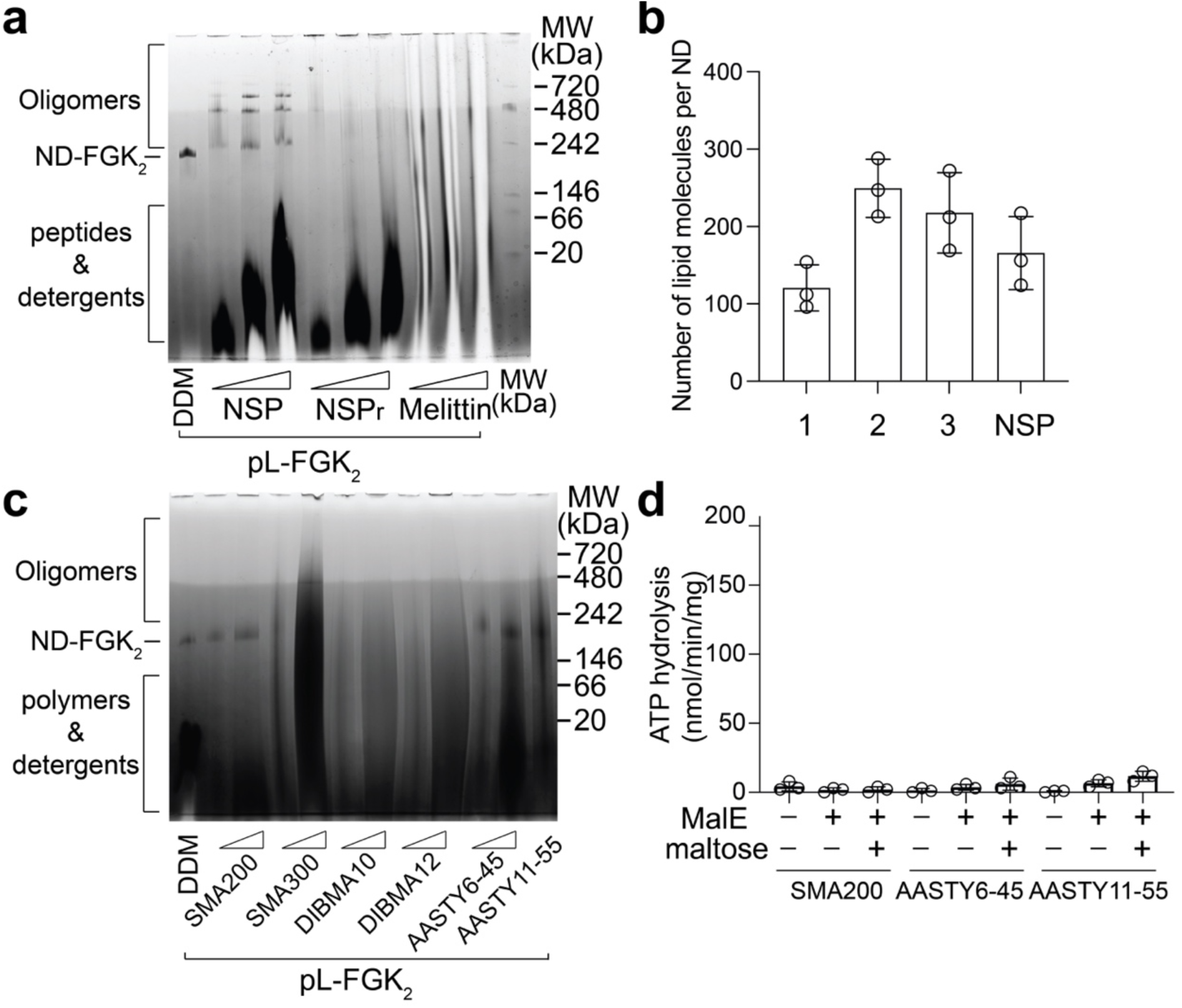
Characterization of MalFGK_2_ NDs extracted from proteoliposomes using peptide and polymer scaffolds. (**a**) Detergent-free ND formation by peptide scaffolds. MalFGK_2_ proteoliposomes were incubated with increasing concentrations (33, 100, and 300 µM) of the indicated scaffold peptides or DDM (0.5%) and analyzed by blue native electrophoresis. (**b**) Lipid quantification in MalFGK_2_ NDs formed with the indicated peptide scaffolds. (**c**) Detergent-free ND formation by polymer scaffolds. MalFGK_2_ proteoliposomes were incubated with increasing concentrations (0.5% and 1.5%) of the indicated scaffold polymers and analyzed by blue native electrophoresis. (**d**) ATPase activities of polymer-encased MalFGK_2_ NDs. Data are shown as mean ± s.d., n = 3 independent experiments.

**Extended Data Figure 2.**
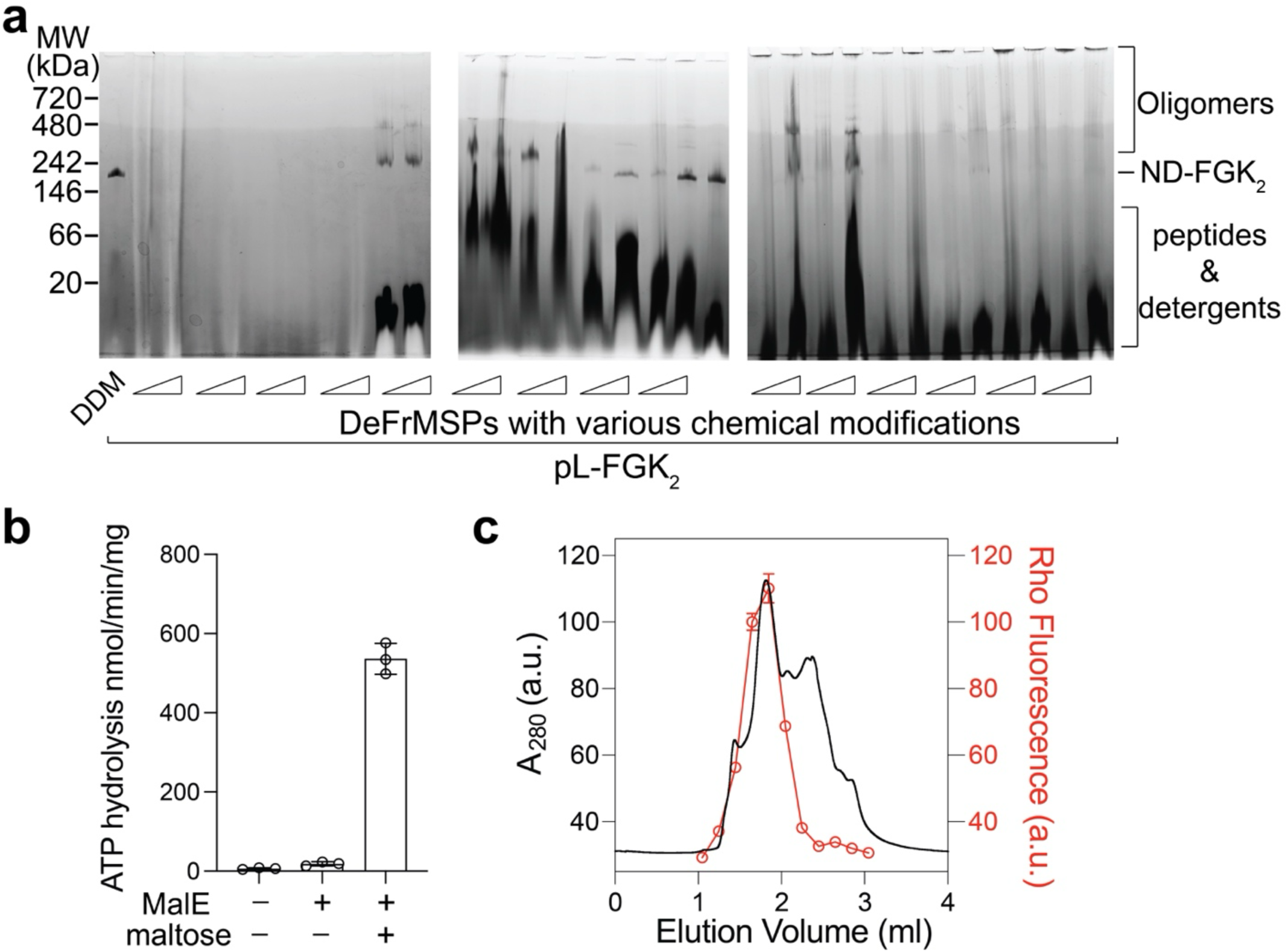
Optimization and characterization of DeFrMSPs for detergent-free formation of MalFGK_2_ NDs. (**a**) Screening DeFrMSP designs to potentiate the performance in extracting MalFGK_2_ from proteoliposomes into NDs. MalFGK_2_ proteoliposomes were incubated with increasing concentrations (33 and 100 µM) of various DeFrMSP designs or DDM (0.5%) and analyzed by blue native electrophoresis. (**b**) ATPase activities of MalFGK_2_ NDs formed with DeFr_5. Data are shown as mean ± s.d., n = 3 independent experiments. (**c**) Lipids were extracted into MalFGK_2_ NDs by DeFr_5. MalFGK_2_ proteoliposomes were prepared with lipid mixtures harboring 0.5% Rho-PE and converted into NDs by DeFr_5. Samples were characterized by SEC using a Superdex200 3.2/300 column.

**Extended Data Figure 3.**
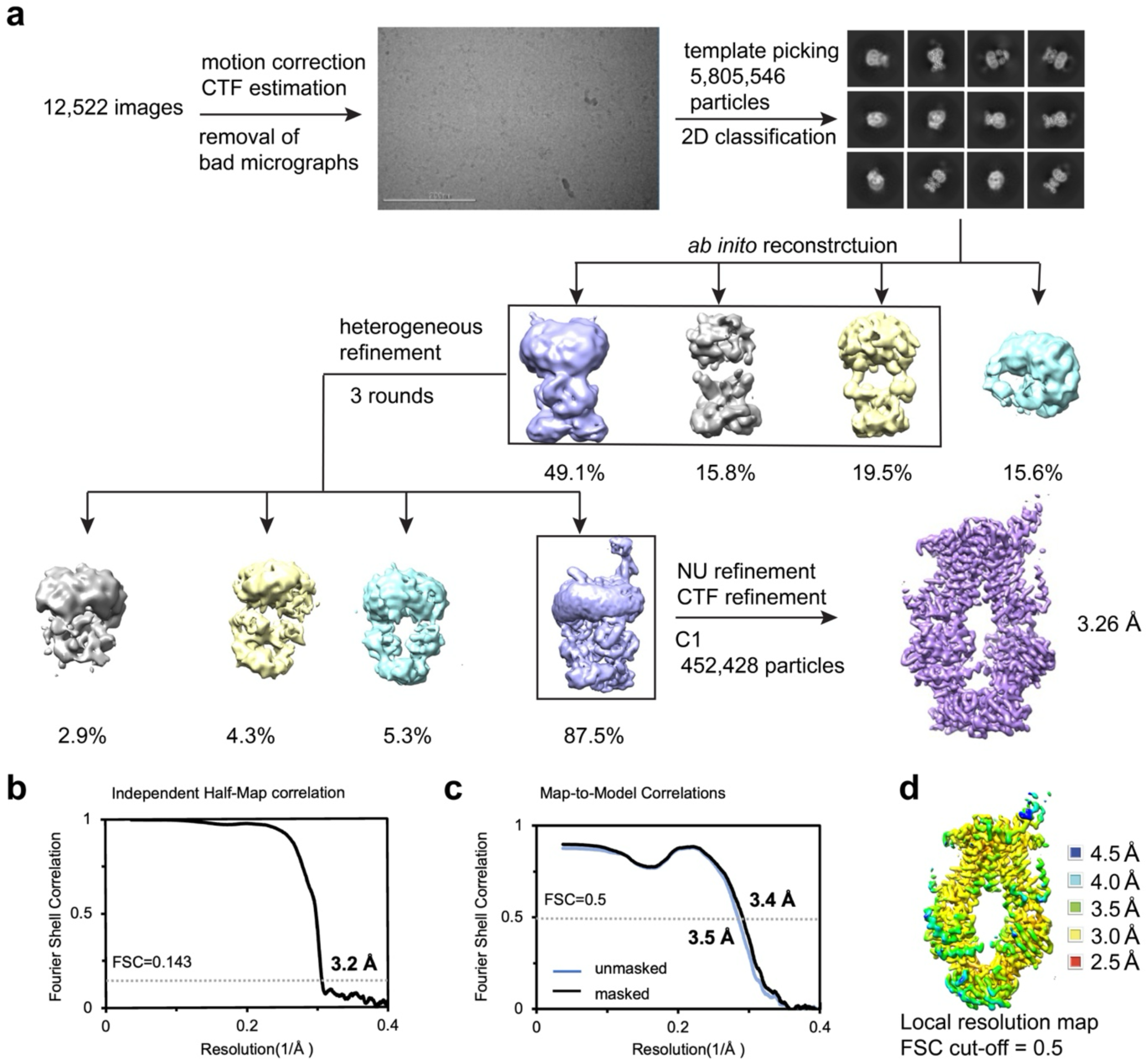
CryoEM analysis of MalFGK_2_ DeFrNDs. (**a**) Overall workflow for data processing. Fourier Shell Correlation (FSC) curves for the overall resolution (**b**), Map-to-model fit between the full map and PDB coordinates (**c**), and local resolution (**d**).

**Extended Data Figure 4.**
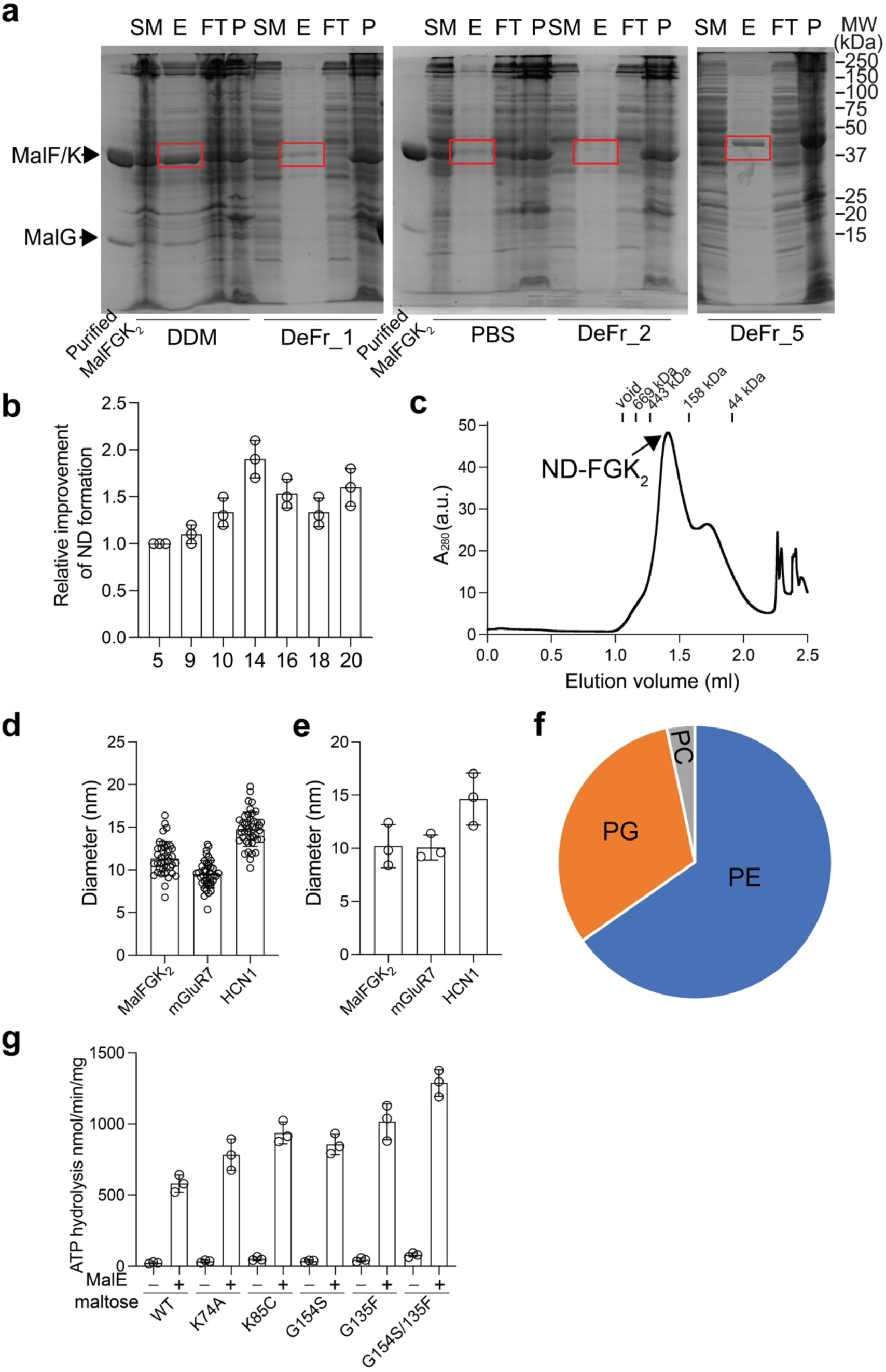
Optimization and characterization of DeFrMSPs for detergent-free extraction of MalFGK_2_ native NDs from crude membranes. (**a**) Crude membranes isolated from cells overexpressing MalFGK_2_ were incubated with the indicated DeFrMSPs or DDM overnight and subjected to pull down experiments. Samples at each step were analyzed by SDS-PAGE. In negative control experiments, PBS buffer with the same amount of DMF was used. Red boxes indicated the extracted MalF and MalK. SM, solubilized materials; E, eluted fractions; FT, flow through fractions; P, insolubilized membrane pellets. (**b**) Evolving DeFrMSP designs based on DeFr_5 for converting MalFGK_2_ proteolipsomes into NDs. Data were quantified based on the density of the MalFGK_2_ complex band normalized to that of the DeFr_5-extracted sample. (**c**) SEC profile of native MalFGK_2_ NDs using a Superdex 200 3.2/300 column. (**d-e**) Diameters of native NDs harboring MalFGK_2_, mGluR7 and HCN1 determined from negative stain EM (d) and DLS (e). (**f**) Lipids associated with native MalFGK_2_ NDs from LC-MS/MS analysis. (**g**) ATPase activities of MalFGK_2_ WT and mutants in native NDs. Data are shown as mean ± s.d., n = 3 independent experiments.

**Extended Data Figure 5.**
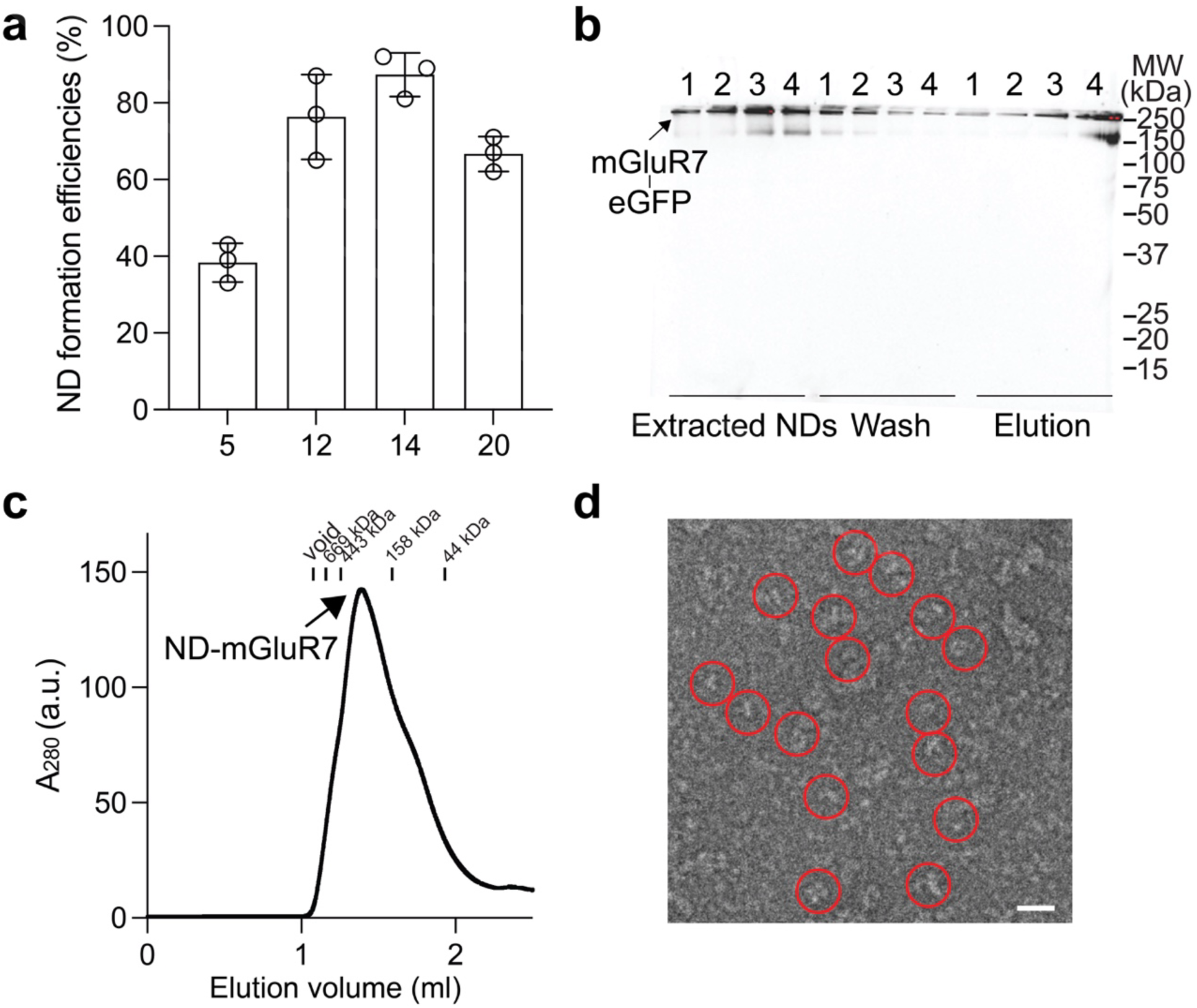
Optimization and characterization of DeFrMSPs for detergent-free extraction of mGluR7 native NDs from crude membranes. (**a**) Screening DeFrMSP designs for extracting mGluR7-eGFP from crude membranes. Solubilized samples were clarified by ultracentrifugation and extracted mGluR7 proteins were quantified by monitoring the fluorescence of eGFP. ND formation efficiencies of mGluR7 were quantified based on the GFP fluorescence emission normalized to that of DDM-solubilized samples. Data are shown as mean ± s.d., n = 3 independent experiments. (**b**) Extracted mGluR7 NDs with different DeFrMSP designs were subjected to pull down experiments. Samples at each step were analyzed by SDS-PAGE and in-gel fluorescence imaging. 1, DeFr_5; 2, DeFr_12; 3, DeFr_14; 4, DeFr_20. (**c**) Fractionation of mGluR7 native NDs using a Superdex 200 3.2/300 column. (**d**) Representative negative stain EM micrograph of mGluR7 native NDs formed by DeFr_14. Scale bar, 30 nm.

**Extended Data Figure 6.**
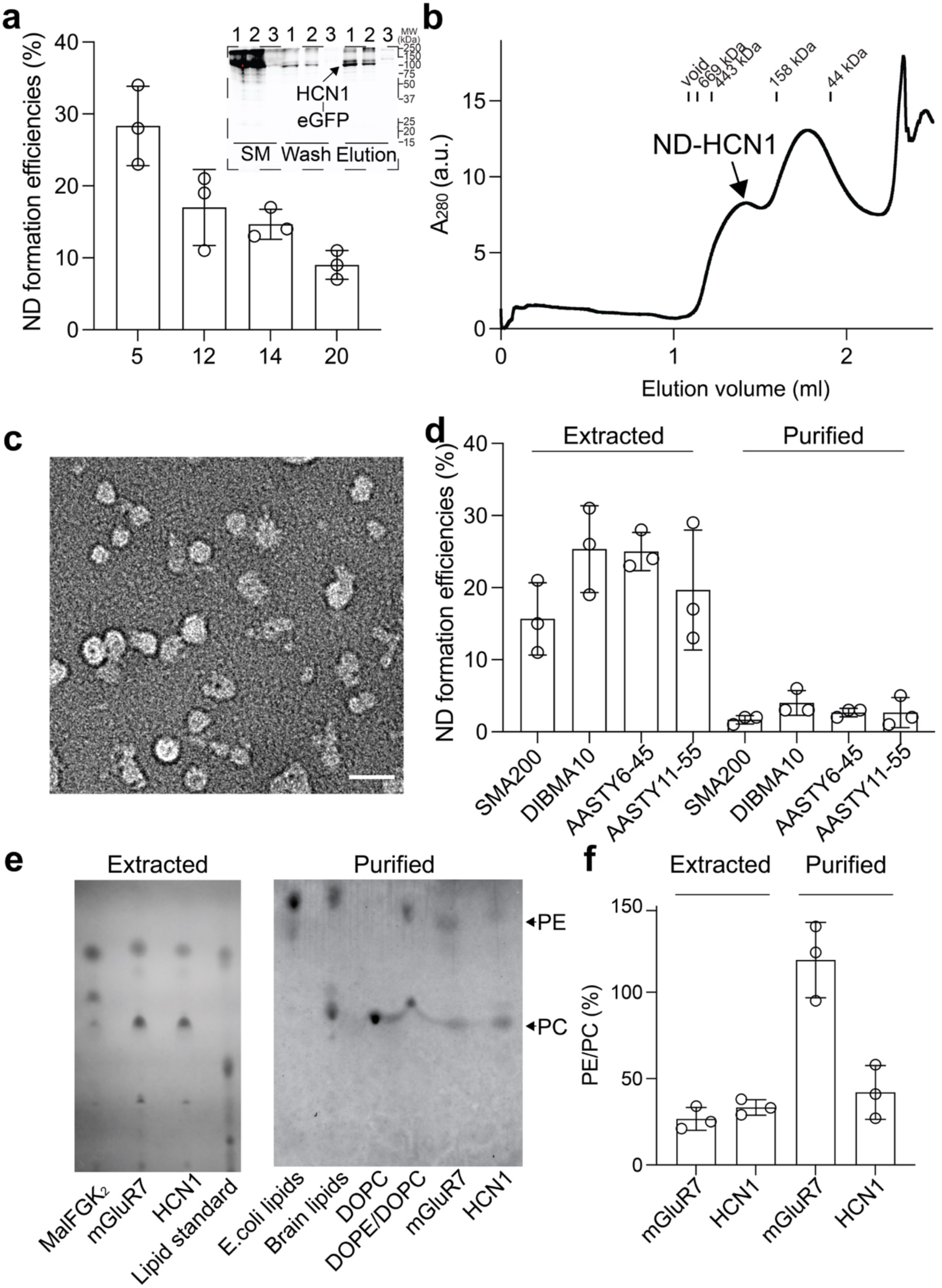
Optimization and characterization of DeFrMSPs for detergent-free extraction of HCN1 native NDs from crude membranes. (**a**) Screening DeFrMSP designs for extracting HCN1-eGFP from crude membranes. Solubilized samples were clarified by ultracentrifugation and ND formation efficiencies of HCN1 were quantified based on the GFP fluorescence emission normalized to that of DDM-solubilized samples. Inset: extracted HCN1 NDs with different DeFrMSP designs and negative control (PBS) were subjected to pull down experiments. Samples at each step were analyzed by SDS-PAGE and in-gel fluorescence imaging. 1, DeFr_5; 2, DeFr_14; 3, PBS. Data are shown as mean ± s.d., n = 3 independent experiments. SM, solubilized materials. (**b**) Fractionation of HCN1 native NDs using a Superdex 200 3.2/300 column. (**c**) Representative negative stain EM micrograph of HCN1 native NDs formed by DeFr_5. Scale bar, 30 nm. (**d**) Extraction of HCN1 native NDs using the indicated polymers. The majority of the extracted proteins could not be purified through affinity purification. Data are shown as mean ± s.d., n = 3 independent experiments. (**e**) TLC analysis of lipids in the indicated native NDs from extracted samples (left) or after purification (right). (**f**) Quantification of PE/PC lipid ratios for the indicated extracted and purified samples. Data are shown as mean ± s.d., n = 3 independent experiments.

**Extended Data Figure 7.**
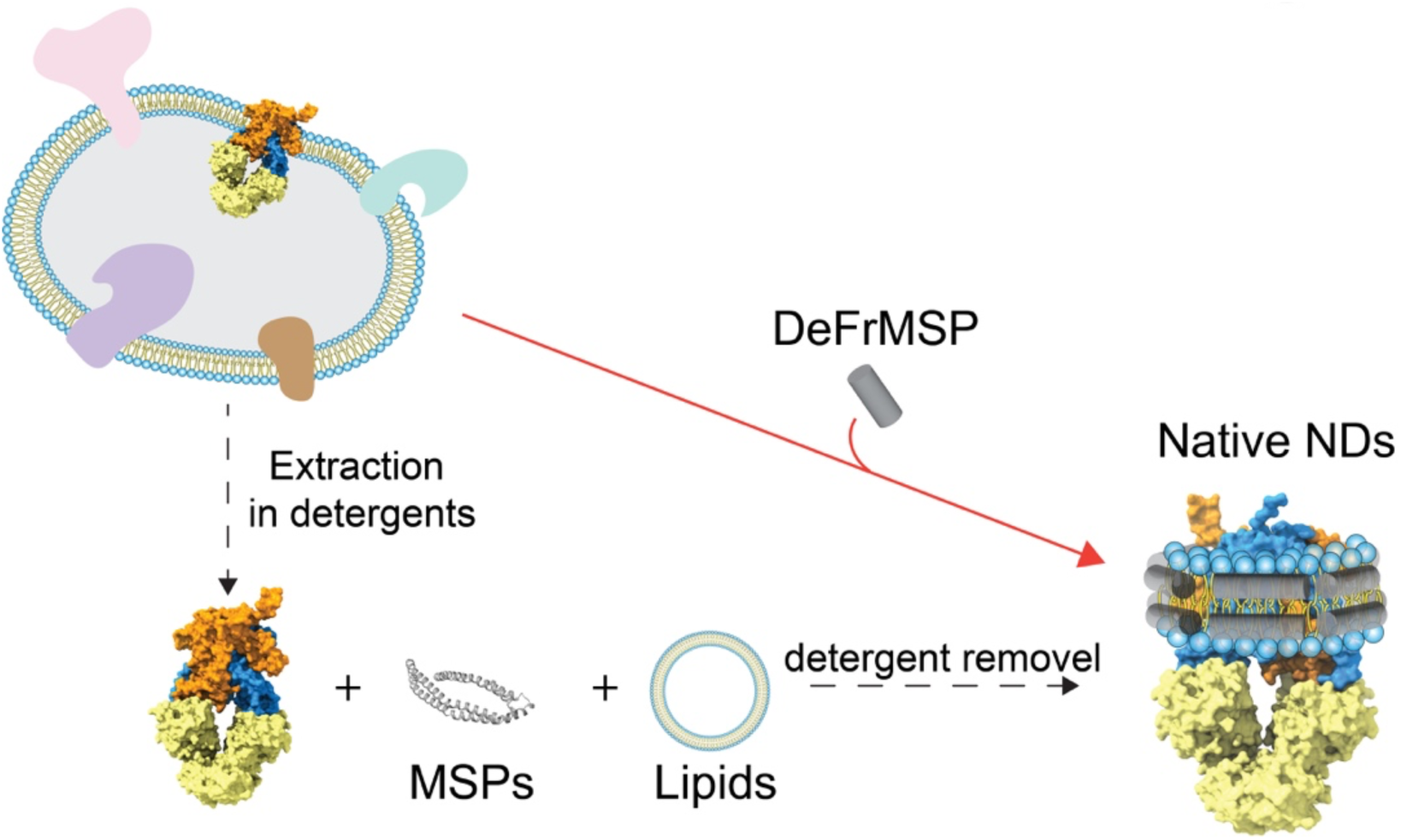
One-step reconstitution of native NDs using DeFrMSPs. Traditional ND reconstitution requires the purification of the membrane protein of interest in detergent micelles (dash line) and optimization of experimental procedures to assemble with MSPs and lipids upon the removal of detergents. In contrast, our approach can directly extract membrane proteins into NDs with native lipids in one step, bypassing the need and limitation of detergent-mediated reconstitution.

